# Dual RNA-seq reveals a type 6 secretion system-dependent blockage of TNF-α signaling and BicA as a *Burkholderia pseudomallei* virulence factor important during gastrointestinal infection

**DOI:** 10.1101/2022.04.22.489234

**Authors:** Javier I. Sanchez-Villamil, Daniel Tapia, Nittaya Khakum, Steven G. Widen, Alfredo G. Torres

**Author notes:** National Polytechnic Institute, National School of Biological Sciences. México city, México. The Ragon Institute of Massachusetts General Hospital, The Massachusetts Institute of Technology and Harvard University, Cambridge, MA. These authors contributed equally to this work (Javier I. Sanchez-Villamil, Daniel Tapia).

## Abstract

Melioidosis is a disease caused by the Gram-negative bacillus *Burkholderia pseudomallei* (*Bpm*), commonly found in soil and water of endemic areas. Naturally acquired human melioidosis infections can result from either exposure through percutaneous inoculation, inhalation, or ingestion of soil-contaminated food or water. Our prior studies recognized *Bpm* as an effective enteric pathogen, capable of establishing acute or chronic gastrointestinal infections following oral inoculation. However, the specific mechanisms and virulence factors involved in the pathogenesis of *Bpm* during intestinal infection are unknown. In our current study, we standardized an *in vitro* intestinal infection model using primary intestinal epithelial cells (IECs) and demonstrated that *Bpm* requires a functional T6SS for full virulence. Further, we performed dual RNA-seq analysis on *Bpm*-infected IECs to evaluate differentially expressed host and bacterial genes in the presence or absence of a T6SS. Our results showed a dysregulation in the TNF-α signaling via NF-κB pathway in the absence of the T6SS, with some of the genes involved in inflammatory processes and cell death also affected. Analysis of the bacterial transcriptome identified virulence factors and regulatory proteins playing a role during infection, with association to the T6SS. By using a *Bpm* transposon mutant library and isogenic mutants, we showed that deletion of the *bicA* gene, encoding a putative T3SS/T6SS regulator, ablated intracellular survival and plaque formation by *Bpm* and impacted survival and virulence when using murine models of acute and chronic gastrointestinal infection. Overall, these results highlight the importance of the type 6 secretion system in the gastrointestinal pathogenesis of *Bpm*.

## Introduction

*Burkholderia pseudomallei* (*Bpm*) is a Gram-negative bacterium and the causative agent of human melioidosis (Galyov, Brett et al. 2010, Wiersinga, Virk et al. 2018). Melioidosis is becoming a health issue due to its increased global awareness and the recognition of the disease as an underreported neglected tropical disease (Savelkoel, Dance et al. 2021). For example, human melioidosis in northeast Thailand ranks as the third most common cause of death due to infectious diseases (after HIV and tuberculosis) (Limmathurotsakul, Wongratanacheewin et al. 2010, Meumann, Cheng et al. 2012), while in other parts of the world, melioidosis is being recognized as endemic in countries located within the tropics (Limmathurotsakul, Golding et al. 2016, Mukhopadhyay, Shaw et al. 2018, Rakotondrasoa, Issack et al. 2018, Rolim, Lima et al. 2018, Sanchez-Villamil and Torres 2018, Smith, Hanson et al. 2018, Steinmetz, Wagner et al. 2018). Worldwide, human and animal infections by this pathogen are significantly underreported, while the number of cases continues to increase (Limmathurotsakul, Golding et al. 2016, Gassiep, Armstrong et al. 2020). A comprehensive epidemiological study estimated that 165,000 melioidosis cases and 89,000 deaths occur every year around the world (Limmathurotsakul, Golding et al. 2016). While clinical manifestations of the disease are diverse, including acute sepsis, chronic localized pathology, or a latent infection that can reactivate decades later. Further disease surveillance is hindered by a limited understanding of the disease and the association of non-specific symptoms that often mimic other persistent infections (Gassiep, Armstrong et al. 2020, Jamani, Mohd Nor et al. 2021). Community acquired cases of melioidosis are likely a consequence of the saprophyte bacterium in contaminated soil or water entering the host through cuts or skin abrasions, ingestion, or inhalation (Limmathurotsakul and Peacock 2011, Wiersinga, Virk et al. 2018). Treatment of the disease is difficult and includes parenterally delivered ceftazidime or carbapenem for 10-14 days, followed by oral trimethoprim-sulfamethoxazole for 12-20 weeks (Estes, Dow et al. 2010, Wiersinga, Virk et al. 2018).

*Bpm* can bind and infect phagocytic and non-phagocytic cell types (Pruksachartvuthi, Aswapokee et al. 1990, Jones, Beveridge et al. 1996) upon coming in contact; however, most of the pathogen-host interactions at the epithelial interface have focused on those infections occurring at the abraded skin or the respiratory mucosal surface, during percutaneous inoculation or respiratory infection (Lazar Adler, Govan et al. 2009, Wiersinga, Virk et al. 2018). For example, attachment to human respiratory epithelial cells was initially thought to be mediated by capsular polysaccharides (Ahmed, Enciso et al. 1999) and type IV pili (Essex-Lopresti, Boddey et al. 2005). However, the function of the capsule in the internalization process has been questioned, and the role of the type IV pili in adherence has only been demonstrated during infection of the respiratory tract (Essex-Lopresti, Boddey et al. 2005, Wiersinga, Virk et al. 2018). In contrast, the exact mechanism of entry and dissemination using other ports of entry remain relatively unexplored and an in-depth analysis of the bacterial mechanisms mediating gastrointestinal (GI) infection have not been undertaken. One prior study investigated whether *Bpm* was able to cause GI infection in mice and it was reported that *Bpm* is capable of infecting intestinal cells following oral inoculation and that chronically colonized GI cells might be the reservoir for dissemination to extra-intestinal sites (Goodyear, Bielefeldt-Ohmann et al. 2017). This has been further validated in non-human primates where it was demonstrated that *Bpm* ingestion serves as the route for disseminated infection (Nelson, Nunez et al. 2021). Due to the intracellular nature of this pathogen, it is critical to investigate the molecular mechanisms mediating GI invasion and dissemination to target organs, and we have previously found that the type 1 fimbriae is involved in initial attachment to the intestinal cells, while the type 6 secretion system participates in cell-to-cell spread (Sanchez-Villamil, Tapia et al. 2020).

Although the exact mechanism of *Bpm* invasion to epithelial cells, particularly intestinal cells, remains unknown, it has been demonstrated that inhibition of actin polymerization reduces the level of entry and invasion (Stone, DeShazer et al. 2014, Wiersinga, Virk et al. 2018). *Bpm* possesses multiple secretion systems, which enable the translocation of proteins into the host cells, favoring invasion and cell-to-cell spread. Rearrangement of the host actin cytoskeleton is induced by effector proteins injected by the type 3 secretion system (T3SS) and autotransporter proteins that mediate actin-based motility (Stevens, Friebel et al. 2003, Lazar Adler, Govan et al. 2009, Lazar Adler, Stevens et al. 2011, Wiersinga, Virk et al. 2018). The T3SS is crucial for vesicle escape, while the autotransporter BimA interacts with monomeric actin in one pole of the bacteria, resulting in formation of actin tails promoting *Bpm* intracellular motility (Stone, DeShazer et al. 2014). Once *Bpm* moves freely around the host cytosol, it approaches neighboring cells, stimulating cell fusion and the formation of multinucleated giant cells (MNGCs) (Whiteley, Meffert et al. 2017). *Bpm* is believed to manipulate host cells by utilizing the type 6 secretion system (T6SS), forming MNGCs for intercellular spread and to avoid interaction with the immune system (Lennings, West et al. 2019). Interestingly, *Bpm* possess five T6SS, which share similarities with other T6SS of Gram-negative pathogens (Miyata, Bachmann et al. 2013). However, only the T6SS-1 (or also known as the T6SS-5) is utilized by *Bpm* to manipulate the host cells for intracellular spread (Lennings, West et al. 2019) and to stimulate the formation of MNGC, being cell fusion the pathogenic hallmark of *Bpm* and of a small group of intracellular bacterial pathogens (Whiteley, Meffert et al. 2017, Lennings, West et al. 2019, Stockton and Torres 2020).

We have previously demonstrated, using a human intestinal epithelial cell line and mouse primary intestinal epithelial cells (IEC), that *Bpm* adheres, invades, and forms MNGCs, ultimately leading to cell toxicity (Sanchez-Villamil, Tapia et al. 2020). Further, we demonstrated that *Bpm* requires a functional T6SS for full virulence, bacterial dissemination, and lethality in mice infected by the intragastric route (Sanchez-Villamil, Tapia et al. 2020). However, several important questions remain unsolved including the mechanism underlying T6SS activity and its physiological role during infection, particularly during GI cell-to-cell spread. Therefore, in this paper we utilized a dual RNA-Seq analysis of infected IECs with wild type (WT) *Bpm* and a type 6 secretion mutant (Δ*hcp1*) to determine which host and pathogen genes are differentially expressed during infection. Several genes up- and down-regulated were identified while comparing the *Bpm* WT and mutant strain and to further validate our analysis, we showed a dysregulation in the TNF-α signaling via NF-κB pathway in the host cells. Further, we demonstrate a regulatory control of the BicA protein, a T3SS, as a potential virulence factor during IG infection by *Bpm*, and an associated virulence factor with bacterial invasion of IECs.

## Results

### Defining intestinal host IECs-*Bpm* transcriptomics using Dual RNA-Seq

To start understanding the molecular mechanism that governs *Bpm* and the host cells during infection of intestinal epithelial cells, we performed a dual RNA-seq analysis to define the impact of expressing the T6SS in the intracellular host environment, using the transcriptome of the pathogen and that of the host cell as readout. Dual RNA-seq has been previously successfully used to profile gene expression simultaneously of an intracellular pathogen and its infected host cell (Westermann and Vogel 2018, Mika-Gospodorz, Giengkam et al. 2020, Penaranda, Chumbler et al. 2021, López-Agudelo, Baena et al. 2022). Therefore, we performed a fully validated dual RNA-seq experiment and then focused our analysis in subsets of bacterial and host proteins differentially expressed during intracellular infection of the *Bpm* wild type K96243 (Titball, Russell et al. 2008) and its Δ*hcp1* mutant (Sanchez-Villamil, Tapia et al. 2020). Briefly, primary intestinal epithelial cells (IECs) derived from rat were selected as host cells and the bacterial infections were performed using *Bpm* WT and *Bpm* Δ*hcp1* at an MOI of 10 during 6 and 12 h of growth (**Fig. 1**, step 1). Total RNA was collected from infected cells (step 2) and pure RNA samples (bacterial or host rRNA was depleted) were used (step 3). The genomic libraries were prepared and sequenced using next-generation sequencing (NGS) technology (step 4) and the sequences were mapped to the rat (step 5) or *Bpm* K96243 (step 6) reference genomes. Finally, we profiled different expression patterns between *Bpm* WT and *Bpm* Δ*hcp1* with high resolution, after infection of IECs. Differences in expression profiles are presented as heatmap (step 7) and volcano plots (**Figs. 2 and 4**).

**Figure 1.**
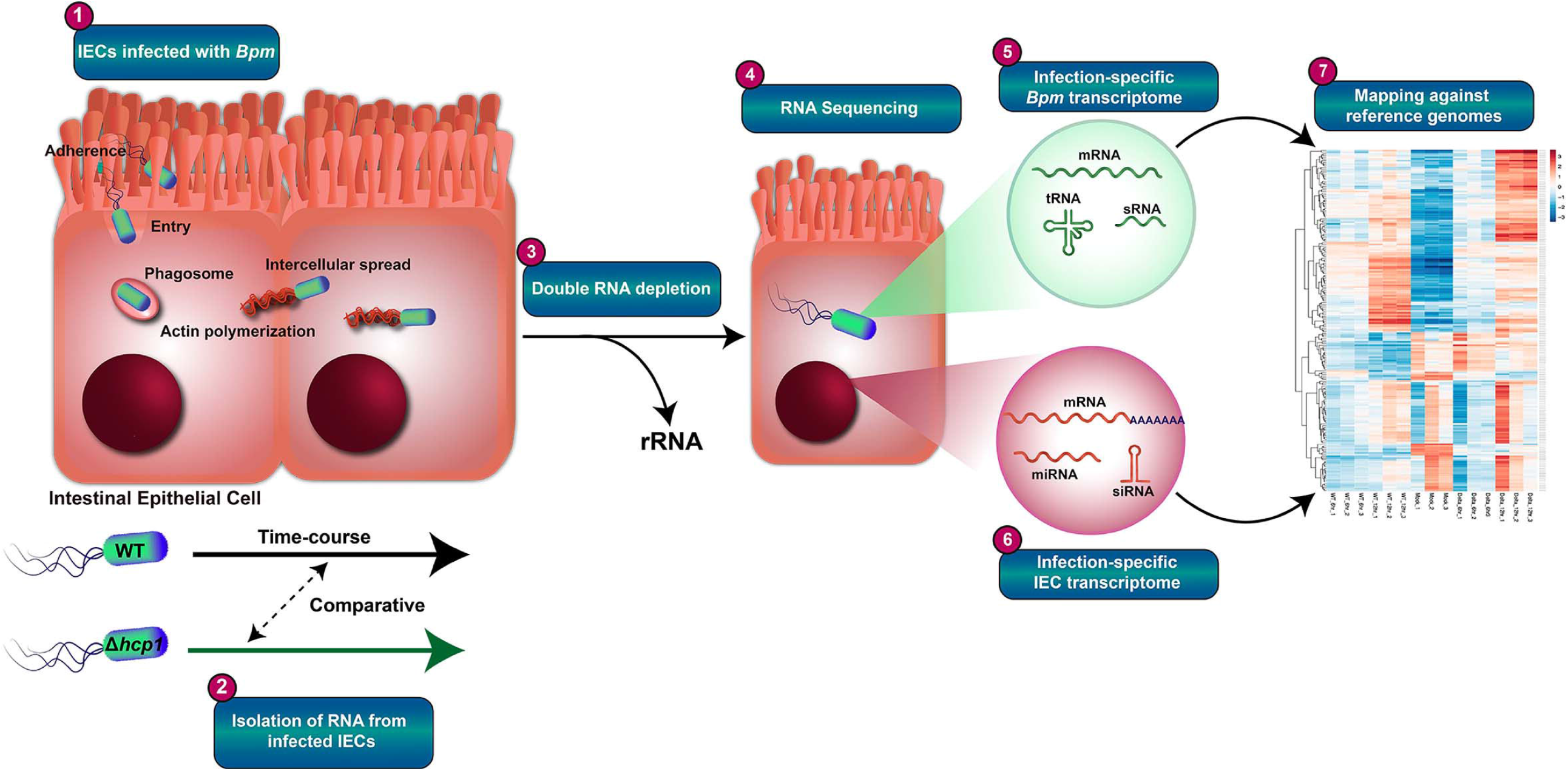
Workflow of dual RNA-seq experiment. High-quality RNA was collected from primary intestinal epithelial cells (IECs) and enriched for either host or bacteria RNA transcripts before creating cDNA libraries. **1**) The IECs were infected with *Bpm* WT K96243 and a *Bpm* Δ*hcp1* mutant for 6 and 12 h using an MOI of 10. **2**) Total RNA was isolated from infected cells and purified RNA samples used for downstream analysis. **3**) Bacterial or host rRNA was depleted using an RNA purification kit. **4**) Purified RNA samples were sequenced using Illumina technology. **5**) Host RNA libraries were mapped against a parent genomic sequence. **6**) Bacterial RNA libraries were mapped against a parent *Bpm* sequence. **7**) Differences in expression profiles were presented as heatmap and volcano plots.

**Figure 2.**
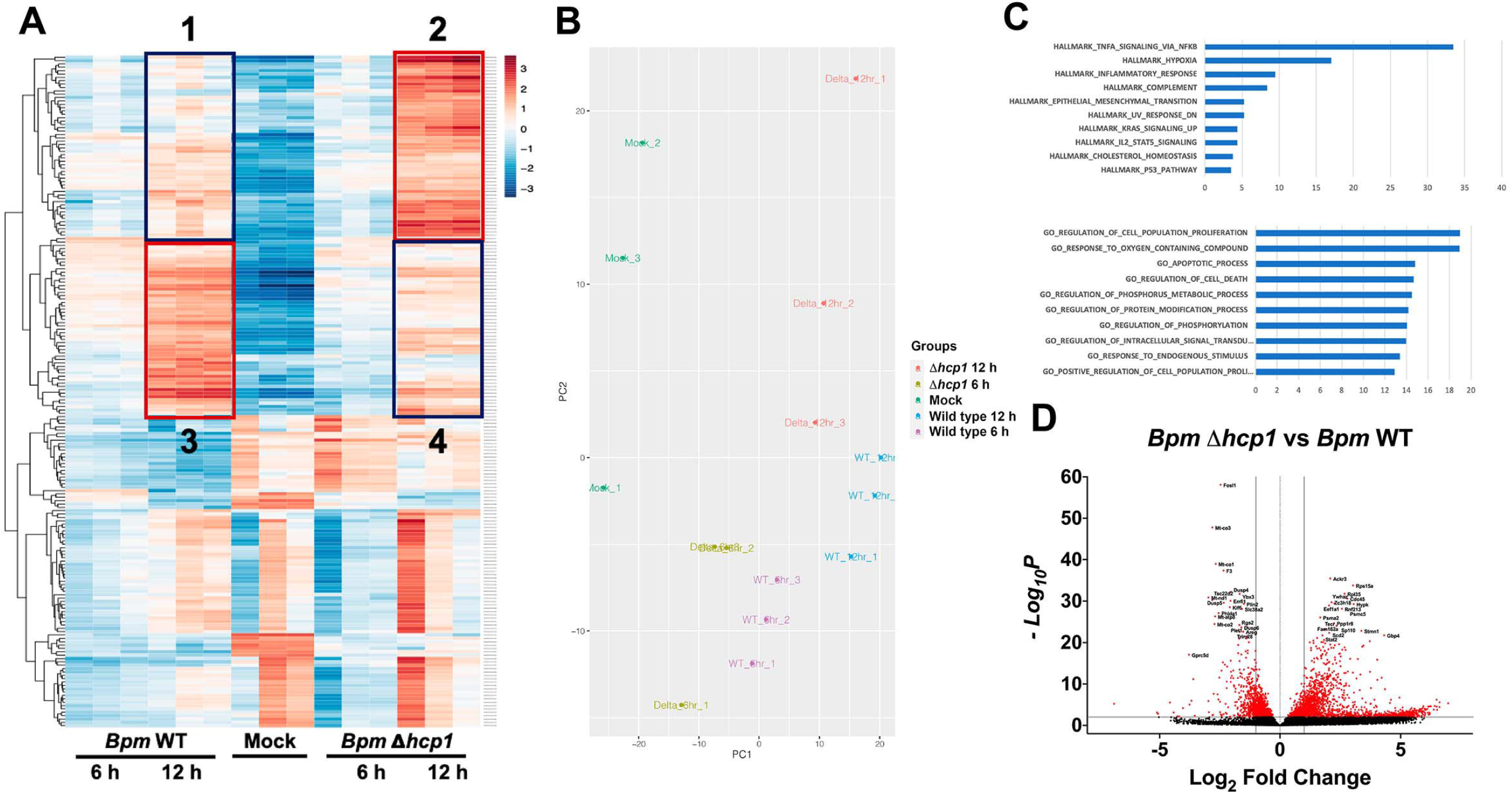
Global host transcriptomic changes in the presence or absence of a T6SS. **A**. Heat map analysis summarizing the transcriptomic differences in the presence or absence of a functional *Bpm* T6SS, showing the 400 most differentially expressed host genes. Genes in box 2 indicated an upregulation in the absence of T6SS as compared to WT in box 1. This suggested a transcriptional repression associated to the T6SS (box 2 vs 1). Genes in panel 4 are downregulated as compared to panel 3, suggesting a transcriptional activation associated to the T6SS **B**. Using ingenuity pathway analysis, hallmark gene association suggests a role for the most differentially expressed genes to be associated with TNF-α signaling via NF-κB. In addition, gene ontology analysis suggests a role of differentially expressed genes linked to cell proliferation, cell death, among other metabolic processes. **C**. The volcano plot demonstrates specific differential expression of genes during *Bpm* IEC infection generated with p-values less than 0.05 in the presence (*Bpm* WT) or absence of a functional T6SS (Δ*hcp1*).

### Differential host gene expression during *Bpm* infection and in the absence of a T6SS

The IECs were infected with *Bpm* WT and *Bpm* Δ*hcp1* for 6 and 12 h and the differential global gene expression were determined (**Fig. 2A**). Panel A shows the host transcriptomic gene differences during infection with *Bpm* in the presence or absence of a functional T6SS at 6 and 12 h post infection. As depicted in the boxes, genes in box 2 are upregulated in the absence of a functional T6SS (Δ*hcp1*) as compared to the WT strain (box 1) and suggesting that a transcriptional gene repression occurs during expression of the T6SS. Conversely, genes in box 4 are downregulated in comparison with those found in box 3, suggesting that a transcriptional activation of those genes occur during expression of the T6SS.

To further establish the association between expression of the T6SS and transcriptional changes in the host cell, 400 of the most differentially expressed genes in IECs were further classified using ingenuity pathway analysis, and we identified an association of several signaling pathways, including those related to TNF-α signaling via NF-κB **(Fig. 2B**). Further, we found that dysregulation of these genes occurs in a T6SS-dependent manner (**Fig. 2C**). Of note, we observed that core responses of *Bpm* interacting with IECs were dominated by a proinflammatory response such as those reported with other cultured epithelial cells (Tan, Chen et al. 2010, Teh, French et al. 2014). We decided to further analyzed these genes because they are found in melioidosis patients and recommended as useful biomarkers of disease progression (Krishnananthasivam, Sathkumara et al. 2017), but also implicate the T6SS-dependent induction of inflammatory response which could be a critical defense mechanisms of IECs against *Bpm*.

### Establishing differential host cell responses to *Bpm* WT and Δ*hcp1* strains

Based on our global differential gene expression analysis, we wanted to further understand specific genes associated with cell signaling pathways affected by the T6SS of *Bpm*. Our volcano plot analysis further confirmed the involvement of the NF-κB signaling pathway (**Fig. 2C**); therefore, we evaluated whether *Bpm* blocks TNF-α-induced NF-κB activation in a T6SS-dependent manner. We first evaluated the biological relevance of NF-κB signaling during *Bpm* infection by imaging NF-κB nuclear translocation in the presence or absence of T6SS (**Fig. 3A**). Our results indicated that *Bpm* sequestered NF-κB in the cytoplasm of IECs, and sequestration is dependent on a functional T6SS, without affecting the intracellular survival of *Bpm*.

**Figure 3.**
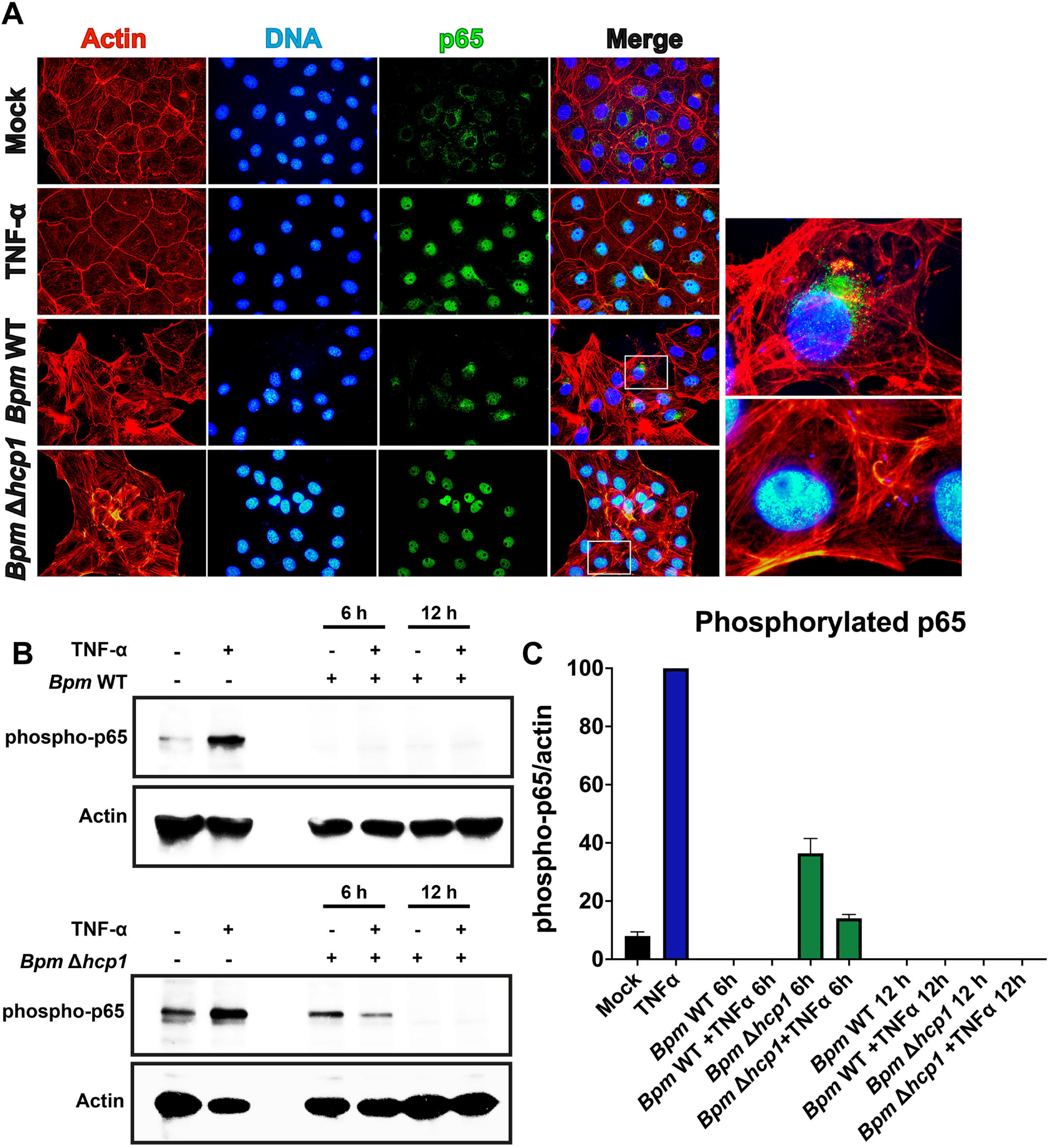
*Bpm* blocks TNF-α-induced NF-κB activation which is dependent on a functional T6SS. **A**. Immunofluorescence of NF-κB nuclear translocation in IECs in the presence or absence of *Bpm* T6SS (Δ*hcp1*), showing sequestration of cytoplasmic NF-κB. The IECs were infected with either *Bpm* WT or *Bpm* Δ*hcp1*. After 12h of infection, cells were fixed, stained, and imaged. Actin cytoskeleton and nuclei were stained with rhodamine phalloidin and DAPI, respectively. **B**. To confirm blockage of NF-κB translocation during *Bpm* infection and dependency of T6SS, the phosphorylation of NF-κB by TNF-α activation was measured by Western blot. Cell lysates from infected IECs were collected after 6 and 12 h of infection with *Bpm* WT or Δ*hcp1*. In addition, separate infected cells were treated for 10 min with TNF-α (20 ng/mL) and subjected to lysis. An anti-phosphorylated p65 antibody was used followed by an HRP-conjugated goat anti-mouse. As a positive control, cells were stimulated with TNF-α only. C. Phosphorylation of p65 was quantified for each infection and presented as a ratio phosphorylation p65/actin.

We then measured the phosphorylation of NF-κB by TNF-α activation by Western blot, where we observed that *Bpm* inhibits NF-κB activation despite TNF-α stimulation post infection (Fig. 3B). We further observed that this inhibition is time dependent and requires a functional T6SS (**Fig. 3B**). Further, we quantified phosphorylation of p65 by densitometry and our results confirmed that this phosphorylation of p65 is partially rescued by 6 h post infection with *Bpm* Δ*hcp1* in the presence or absence of TNF-α (**Fig. 3C**). These results indicate a NF-κB blockage by the T6SS of *Bpm* during late IEC infection. It is well recognized that various virulence factors activated by bacteria during epithelial infection are targets of NF-κB, and often recruit inflammatory cells to facilitate dissemination (Krakauer 2019). In melioidosis infections, it is known that TNF-α and some other interleukins are involved in the early inflammatory response, and NF-κB is the key transcription factor that modulates their expression (Wiersinga, Virk et al. 2018). These results suggest a role of the T6SS system in controlling NF-κB blockage, and it is dependent on additional factors that may be related to early infection events, such as the T3SS. Together, our results, along with other studies, (Tan, Chen et al. 2010) confirmed that *Bpm* prevents the initiation of the NF-κB, thus reducing host inflammatory responses by utilizing T6SS proteins.

### Defining *Bpm* transcriptomics during IECs infection

We used principal component analysis (PCA) plots of the dual RNA-Seq data comparing *Bpm* WT and *Bpm* Δ*hcp1* to show that different conditions cluster together (data not shown). Then, a volcano plot of the 200 most differentially expressed bacterial genes associated with *Bpm* IEC infection was generated (genes with *p*-value <0.05; **Fig. 4A**). We observed a dysregulation in gene expression associated with two important virulence mechanisms namely, the T3SS, associated genes (*bopC, bicA, bprD, bopA, bpspV*) and the T6SS *(tssD, tssM, tssG, tagL-2)* (**Fig. 4A**) (Sun and Gan 2010). To determine whether the gene products participate in *Bpm* intracellular survival or cell-to-cell spread, we used a *Bpm* transposon mutant library (library comprised of a total of 8,729 unique transposon mutants in the *Bpm* 1026b strain) and evaluated several of the mutants for their ability to form plaques (**Fig. 4B**). We included some transposon mutants associated with the T3SS (*bpspV, bopC, bopA, bicA*), T6SS (*tagL-2*) and other genes (*bhuR, hfq*, etc.) and determined whether they affected plaque formation. We confirmed that a T6SS (Tn::*tagL-2*) mutant was unable to form plaques, but also found that some T3SS mutants reduced plaque formation (*Fig. 4B*). Of notice, we identified one protein encoded by the *bicA* gene that impacted cell-to-cell spread. The BicA protein forms a complex with BsaN to regulate the expression of T3SS effectors, but it has also been proposed as a regulator affecting the expression of the T6SS (Sun and Gan 2010). To corroborate our results, we wanted to further investigate the function of BicA as a virulence factor during IECs infection.

**Figure 4.**
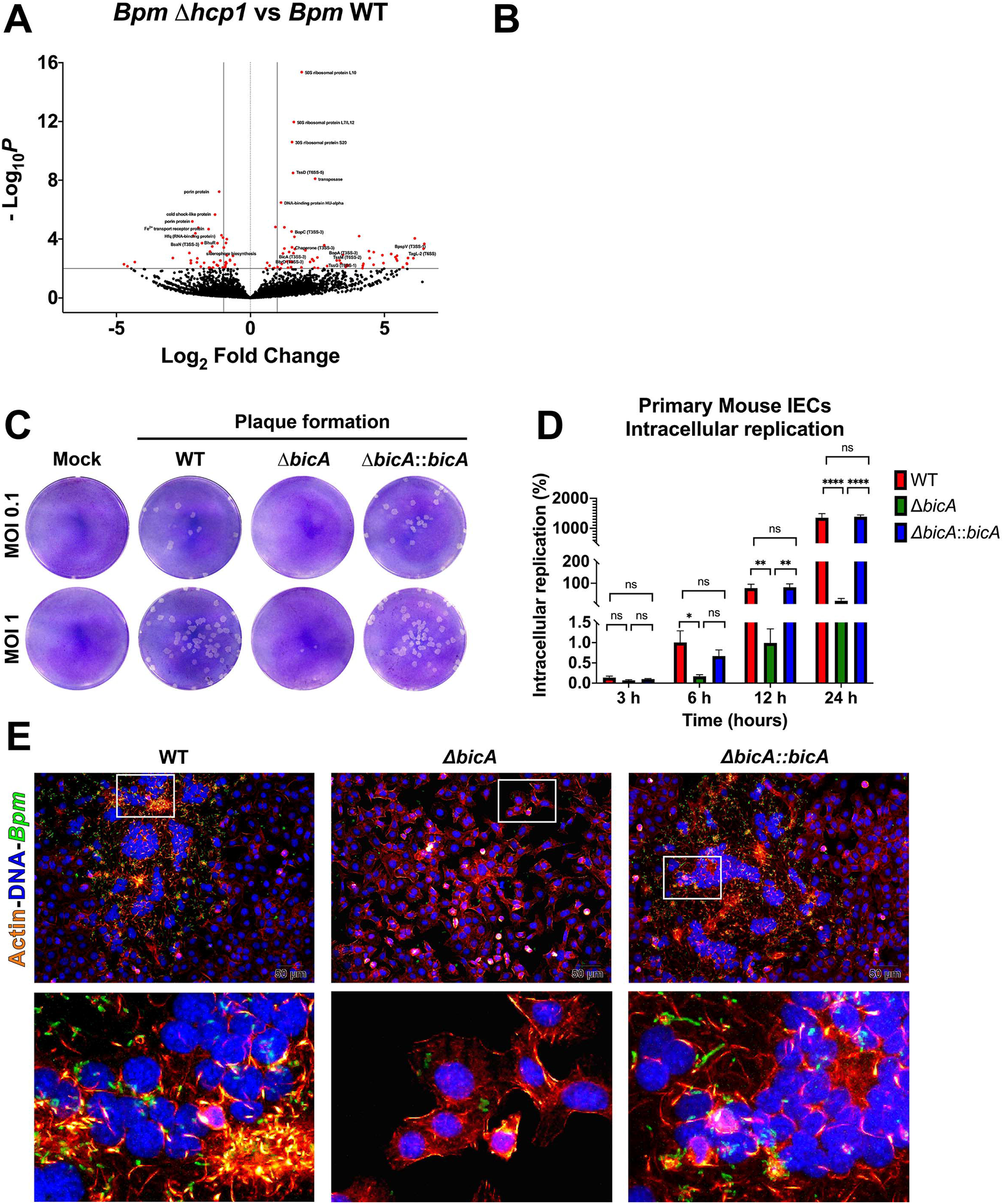
Differential expression profile of *Bpm* in the absence of a T6SS, reveals a role for BicA as a potential virulence factor associated with IEC infection. **A**. Volcano plot of the 200 most differentially expressed genes associated with *Bpm* IEC infection generated with genes with a p-values less than 0.05. The plot shows dysregulation of gene expression associated with two important mechanisms of virulence, namely, T3SS and T6SS. **B**. Using a transposon library, several mutants dysregulated during IEC infection were tested for their ability to form plaques. **C**. A Δ*bicA* mutant and a complemented Δ*bicA*::*bicA* were generated and tested for plaque assays (MOI 0.1 or 1) at 24 h. **D**. IECs were infected with *Bpm* WT, Δ*bicA* or Δ*bicA*::*bicA* for 1 h using an MOI of 10. After infection, extracellular bacteria were killed using 1 mg/mL of kanamycin for 1 h. Intracellular bacterial replication was quantified by CFU enumeration at 3, 6, 12 and 24 h. Error bars of the mean represent the average ± standard error of two experiments with three biological replicates. Statistical analysis was done using a one-way ANOVA followed by a Sidak multiple comparison test. \**p* < 0.05, \*\**p* < 0.01, \*\*\*\**p*⍰<⍰0.0001, ns, not significant. **E**. Primary IECs were infected with *Bpm* WT, Δ*bicA* or Δ*bicA*::*bicA* for 6 or 12 h at 37°C and 5% CO_2_, using an MOI of 10. After infection, cells were washed with PBS, fixed, and stained, prior to microscopic examination (20×). Bacterial cells were stained with sera collected from mice that immunized with *Bpm* PBK001 live attenuated vaccine, followed by an Alexa fluorophore 488 goat anti-mouse IgG, IgM, IgA (H+L) secondary antibody. Actin was stained using rhodamine-phalloidin and cell nuclei with DAPI. Magnified views (×5) are shown on the bottom panels. Images were processed and analyzed using ImageJ analysis software.

### Defining the function of BicA as a virulence factor during IEC infection

To validate our global gene expression analysis, we focus our attention in one protein that is proposed as a chaperone/regulator interacting with BsaN, which is known to participate in the regulation of the T3SS expression, but has also been associated with the expression of T6SS genes (Sun and Gan 2010). Therefore, we constructed a *Bpm* Δ*bicA* mutant and Δ*bicA::bicA* complemented strains and confirmed the plaque phenotype observed with the *bicA*::*Tn* (*Tn*::1629) mutant, at two different MOIs (**Fig. 4C**). Our results showed that the Δ*bicA* mutant was unable to form plaques, a phenotype reversible by complementation. Then, we also evaluated whether the Δ*bicA* mutant had an impact on intracellular replication. The IECs were infected with the *Bpm* strains for 1 h, followed by kanamycin treatment to kill extracellular bacteria and then incubated at different time points. Our results at ^3, 6, 12^ and 24 h post-infection demonstrated that the Δ*bicA* mutant displayed a survival defect in the intracellular compartment and that a functional *bicA* gene restores the ability of *Bpm* to survive intracellularly (**Fig. 4D**). Finally, we used immunofluorescence analysis to visualize the intracellular replication and dissemination of the different *Bpm* strains. We showed that the *Bpm* WT and complemented strains invade, disseminate, and spread from cell-to-cell, inducing the formation of MNGCs (**Fig. 4E**). In contrast, the Δ*bicA* mutant was unable to disseminate, spread from cell-to-cell or mediate MNGC formation (**Fig. 4E**). Overall, we have validated the dual-RNA-Seq analysis and demonstrate that BicA plays a role in controlling T6SS genes, resulting in the inability of the Δ*bicA* mutant to disseminate within IECs.

### *Bpm* Δ*bicA* mutant can colonize the GI tract but is unable to disseminate to target organs in acute and chronic mouse models of infection

We have previously developed and optimize murine models of acute and chronic melioidosis infection (Sanchez-Villamil, Tapia et al. 2020) and used to assess virulence of *Bpm* WT, Δ*bicA* or Δ*bicA*::*bicA* strains when delivered by the intragastric route. To validate the significance of BicA, as well as the importance of the pathways identified by our RNA seq analysis on rat cells, we performed *in vivo* animal mouse infection experiments. Cells used for the RNA sequencing experiment were commercially acquired as mouse primary IECs; however, the sequencing results found that the DNA reads were closer associated with the rat genome. For the acute GI murine model, we infected animals with 2.5 LD_50_ of *Bpm* WT, Δ*bicA* or Δ*bicA*::*bicA* strains and evaluated survival for 21 days (**Fig. 5A**). Three days after infection, animals infected with *Bpm* WT showed 75% survival and consistent weight loss (**Fig. 5B**) that was recovered as the infection progress. In contrast, 100% survival was observed with the Δ*bicA* or Δ*bicA*::*bicA* strains (**Fig. 5B**). To assess colonization of the GI tract or dissemination by the different *Bpm* strains, we collected segments of the GI tract (stomach, small intestine, colon, cecum) and spleen to quantify bacterial loads, respectively (**Fig. 5C**). Animals infected with *Bpm* WT had recoverable bacteria from stomach, colon, cecum, and spleen, with a limit of detection of 1 CFU. We were unable to recover the Δ*bicA* mutant from stomach or small intestine, and only a few bacteria were recovered from the spleen of infected animals (**Fig. 5C**). Interestingly, the Δ*bicA* mutant as well as the *Bpm* WT strain colonized the colon and cecum. Complementation of the Δ*bicA* mutant restored colonization to wild type levels.

**Figure 5.**
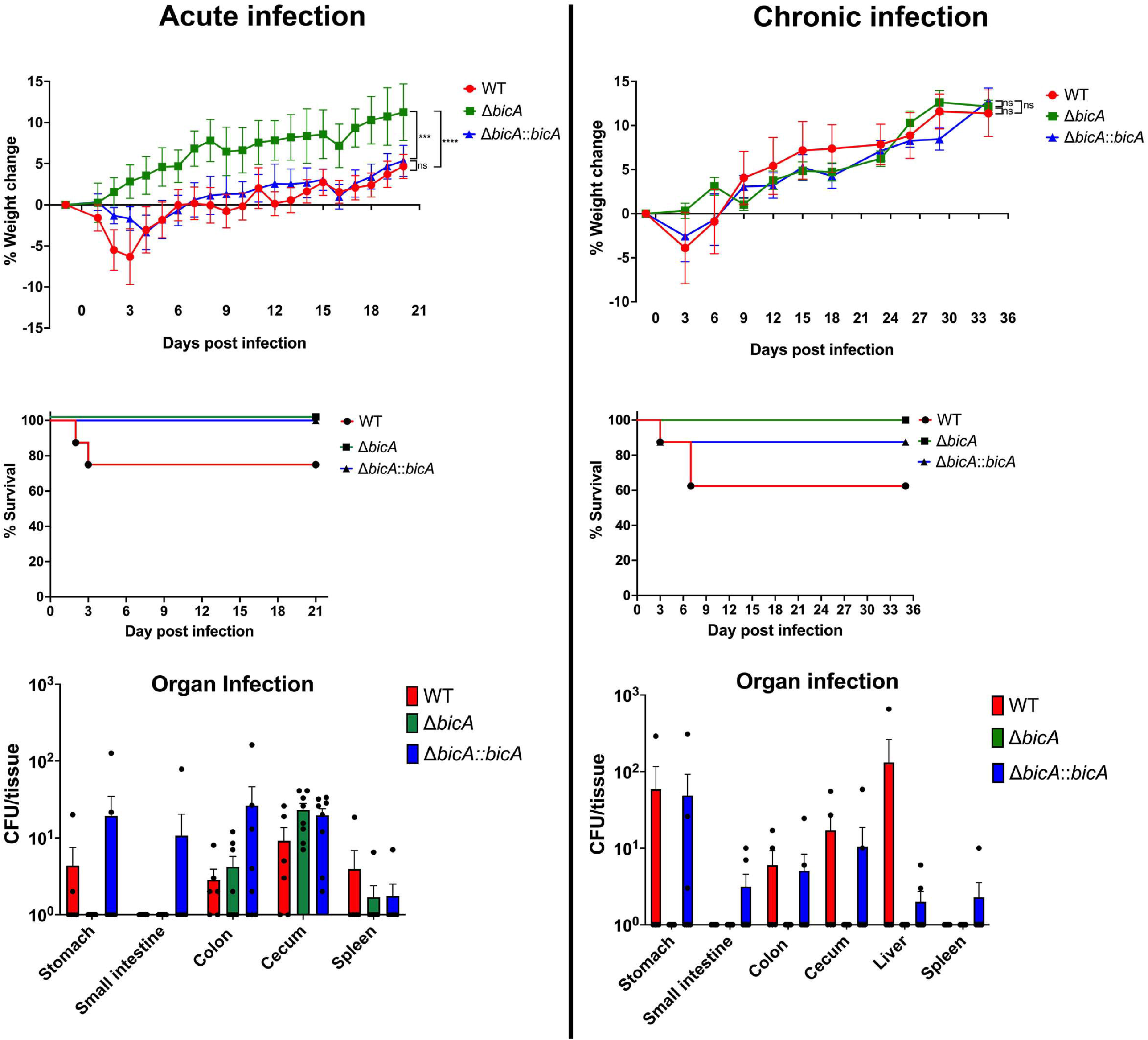
Functional BicA is required for full pathogenesis of *Bpm* in acute and chronic gastrointestinal mouse models of melioidosis infection. Animals (n = 8/group) were infected via gavage with a high-(**A - C**. acute) or low-(**D - F**. chronic) dose equivalents of 2.5 LD_50_ (∼7.5 × 10^6^ CFU/mouse) or 1 LD_50_ (2.5×10^6^ CFU/mouse), respectively, of *Bpm* WT, Δ*bicA* or Δ*bicA*::*bicA*. **A and D**. Weight changes of the infected animals were evaluated for 21 or 35 days, respectively. **B** and **E**. Survival curves of animals infected with each strain in either the acute or chronic infection models. **C** and **F**. Bacterial loads in gastrointestinal tract (stomach, small intestine, colon, and cecum), liver and spleen of the acute and chronic studies were determined for CFU per tissue. Error bars of the mean represent the average +/- standard error.

We further assess whether the *Bpm* strains disseminate to distal organs during a longer infection course using a chronic melioidosis GI infection model. Animals were inoculated with a sublethal dose (∼1 LD_50_) of *Bpm* WT, Δ*bicA* or Δ*bicA*::*bicA* strains and survival was assessed for 35 days (**Fig. 5D**). Animals inoculated with *Bpm* WT or complemented Δ*bicA*::*bicA* strains loss some weight early during infection, which was recovered with time (**Fig. 5D**). Mice inoculated with *Bpm* WT and Δ*bicA*::*bicA* strains exhibited 62.5% or 87.5% survival, respectively, at 35 days post-infection (dpi), while 100% of animals that were inoculated with *Bpm* Δ*bicA* mutant survived to the end of the study (**Fig. 5E**). Bacterial load was evaluated by collecting individual sections of the GI tract as well as the liver and spleen which were processed for CFU enumeration (**Fig. 5F**). Animals inoculated with *Bpm* WT or Δ*bicA*::*bicA* complemented strain exhibited persistent colonization of the stomach, colon, and cecum, while the Δ*bicA*::*bicA* strain was also able to persist in the small intestine (**Fig. 5F**). We were unable to recover the Δ*bicA* mutant strain in any of the intestinal locations tested. *Bpm* WT was able to disseminate and replicate in the liver while the Δ*bicA*::*bicA* strain was also recovered in the liver and spleen (**Fig. 5F**). Like the GI tract segments evaluated, the Δ*bicA* mutant was not recovered neither from the liver nor from the spleen of any infected group (Fig. 5F). Overall, our data demonstrate the ability of *Bpm* to colonize the GI tract and disseminate to target organs and this bacterium requires of a functional *bicA* gene to exert its full virulence phenotype.

## Discussion

Although some progress has been done understanding the pathogenesis of this remarkable intracellular bacterium, not much progress has been done deciphering the host-pathogen interactions that occur during *Bpm* infection, and no information is available about transcriptional changes occurring during the intracellular lifestyle of *Bpm*. We have recently shown the importance of gastrointestinal melioidosis as an understudied route of infection and identified some *Bpm* virulence factors participating in the GI infection and pathogenic process of this intracellular pathogen (Sanchez-Villamil, Tapia et al. 2020). Now, using dual RNA-seq, we were able to identify host and bacterial genes that are differentially expressed during *Bpm* infection and dissemination within gastrointestinal epithelial cells. We were able to identify a handful of host genes that primarily encode for proteins involved in the inflammatory response to infection, but also gene products that mediate *Bpm* regulatory control of secretion systems involved in bacteria entry to the host cell as well as those involved in cell-to-cell spread and eventually MNGS formation. Although the commercially available cells purchased for the RNA seq experiment were acquired as mouse primary IECs, our sequencing results identified closer sequence similarities to the rat genome. However, the cells used in this analysis also demonstrate a consistent phenotype *in vitro* in their ability to permit rapidly multiplying intracellular bacteria and induction of MNGC formation. Therefore, these results allowed us to identify BicA as an important virulence factor in the pathogenesis of *Bpm*.

As presented, the GI tract remains an underreported and understudied point of entry for *Bpm*, particularly in cases of unexplained origin of the disease (Limmathurotsakul, Golding et al. 2016, Goodyear, Bielefeldt-Ohmann et al. 2017). However, recent evidence in non-human primates demonstrated that indeed, ingestion serves as a route for disseminated infection (Nelson, Nunez et al. 2021). Our laboratory has made significant progress in the development of *in vitro* and *in vivo* GI pathogenesis murine models of *Bpm* infection and have identified the Type 6 Secretion System (T6SS) as one of the virulence factors used during GI infection (Sanchez-Villamil, Tapia et al. 2020). We now offer evidence indicating that the T6SS is linked to the host inflammatory response which is dampened during *Bpm* intestinal infection. Although it is unclear whether the immune suppressive effect is true across different *Bpm* strains, our observation indicating that this may be true given that this virulence mechanisms are found in different *Bpm* strains.

A previous study demonstrated that NF-κB activation during *Bpm* infection occurs in a Toll-like (TLR) receptor and MyD88 independent manner which requires the T3SS gene cluster 3 (T3SS3) effector proteins (Teh, French et al. 2014). However, the study was not able to demonstrate that the T3SS directly activates NF-κB, but the authors found that this system facilitates bacterial escape into the cytosol where the host is able to sense the presence of the pathogen leading to NF-κB activation. Further, another study using *Bpm*-infected primary human macrophages implicated the T3SS-3 dependent inflammasome activation and IFN-γ induced immune mechanisms as a defense mechanism against the pathogen, although reduced intracellular *Bpm* loads were observed, the inflammatory response was not completely abolished (Lichtenegger, Stiehler et al. 2020). Now, we present evidence that a functional T6SS is required to block NF-κB activation during *Bpm* infection of IECs and this system seems to work in conjunction with the T3SS to modulate the inflammatory response observed during infection of host cells. One intriguing future direction would be to understand the role BicA plays in T6SS function or a regulatory mechanism controlling the T3SS and T6SS. Furthermore, understanding this mechanism would provide a deeper understanding controlling how the pathogen can disseminate from cell to cell before activation of an effective inflammatory response, allowing the pathogen to reach other target organs.

As demonstrated here and elsewhere, *Bpm* survives and replicates inside host phagocytic and non-phagocytic cells (Wiersinga, Virk et al. 2018, Sanchez-Villamil, Tapia et al. 2020) and the mechanisms by which bacterial virulence factors, particularly the T6SS, interfere with host cell signaling are largely unknown. A significant gap in knowledge exists regarding the bacterial regulatory mechanisms that participate in *Bpm* intracellular survival, cell-to-cell spread, and MNGC formation in epithelial cells. The completion of the dual RNA-seq analysis provided us with the opportunity to understand the molecular consequences of expressing the T6SS in the intracellular host environment. Further, we were able to identify a mutant *Bpm* strain in the *bicA* gene that was defective in plaque formation. The BicA protein has been proposed to form a complex with the regulator BsaN, controlling the expression of genes encoding the T3SS-3, but they have also been proposed as regulators affecting the expression of the T6SS (Sun and Gan 2010). In this study, we wanted to understand the function of differentially regulated genes during *Bpm* infection in the context of controlling the T6SS, its role on cell-to-cell spread, and MNGCs formation; therefore, we further analyzed BicA and its role affecting T6SS-mediated intracellular virulence.

As any other large transcriptomic analysis that tries to elucidate host-bacterial interactions, our study has some limitations. The first difficulty to interpret our data is that we cannot distinguish between differential host responses due to *Bpm* disseminating within the cell cytoplasm and those spreading from cell to cell. However, inclusion of the Δ*hcp1* mutant and subsequent construction of a Δ*bicA* mutant, allowed us to start dissecting the host responses elicited by the bacteria that gets trap within the intracellular environment versus the WT strain that quickly hampered the inflammatory response to escape and disseminate to other organs. A second limitation in the interpretation of our data is that the *Bpm* WT replicates more effectively in the intracellular space than the Δ*hcp1* mutant, producing higher number of reads analyzed in the RNA-Seq analysis. As a result of this difference, we spent time normalizing the different samples and validating our findings with other *in vitro* and *in vivo* experiments to confirm the differences in the virulence phenotypes. Finally, our *Bpm* infection of IECs was relatively high using an MOI 10, which is significantly higher than a natural infection, but this level of infection guaranty that the differences that we are observing between the *Bpm* WT and Δ*hcp1* mutant are identified by the RNA-seq experiment. As demonstrated in our manuscript, the differential expression of host or bacteria genes were further validated with more in-depth experiments that are helping us to have a better picture about the interactions occurring during intracellular survival of *Bpm* in IECs.

The advantage of targeting BicA as the putative regulator, that has been implicated in control of the T6SS, was to provide us with a better understanding of the genes identified in the dual RNA-Seq but also to establish possible mechanism that *Bpm* is exerting to control T3SS and T6SS expression and which occurs during intracellular conditions in the intestinal epithelial cells. To start defining the role of BicA in *Bpm* virulence, we constructed a Δ*bicA* in the *Bpm* K96243 background and the initial characterization indicated that this putative regulator represses T6SS genes during host cell invasion. The BicA-mediated regulation of the T6SS was rescued by complementation of the Δ*bicA* mutant to mimic wild-type levels. Future studies will investigate the details of the regulatory cascade controlling T6SS expression and how this secretion system interacts with the host immune mechanisms, for instance by screening different *Bpm* strains and applying genome- or transcriptome-wide association studies on the bacterial side, or through genetic manipulation or the application of immunomodulating agents on the host side. Our results about the *Bpm* pathogenesis *in vivo* demonstrate the ability of this pathogen to cause disease in animals infected via the oral route. Furthermore, the role of BicA controlling the T3SS during early stages of infection is recapitulated in our acute infection model. Although we did not see the same effect in a chronic infection model, this suggest that other virulence factors, or a well-orchestrated regulatory mechanism controlling secretion systems takes place during the different stages of infection. BicA has been demonstrated to activate the T6SS-associated proteins VirA and VirG, which are both important for the polymerization of actin, thus important for bacterial intercellular motility (Chen, Schröder et al. 2014).

In summary, we have used dual RNA-seq to gain insights into the transcriptome structure and mechanisms of gene regulation used by the complex intracellular pathogen, *Bpm* during GI infection. We provide evidence for the strict regulatory control of bacterial genes whose products are associated in hampering the host inflammatory response. We confirmed a relationship between inhibition of NF-κB activation via TNF-α and T6SS expression. These findings are opening the door to further studies on the regulation of gene expression in *Bpm* and its mechanisms of pathogenesis. In general, our study is only one of very few studies characterizing the host-pathogen transcriptome of intracellular bacteria that can inform and provide insights into to the design of novel therapeutics and vaccines that can prevent melioidosis disease.

## Material and Methods

### Bacterial strain and growth conditions

All bacterial strains used in this study are listed in Table 1. *B. pseudomallei* wild-type (1026b or K96243), mutant, and complemented strains were routinely grown in LB agar plates and broth at 37°C. *E. coli* NEB-5α and S17-1 λ*pir* were grown in LB agar plates and broth at 37°C, and when appropriate Kanamycin (50 or 100 µg/ml) was added for plasmid selection. For the counter selection, co-integrants were grown in YT medium supplemented with 15% sucrose.

**Table 1.**
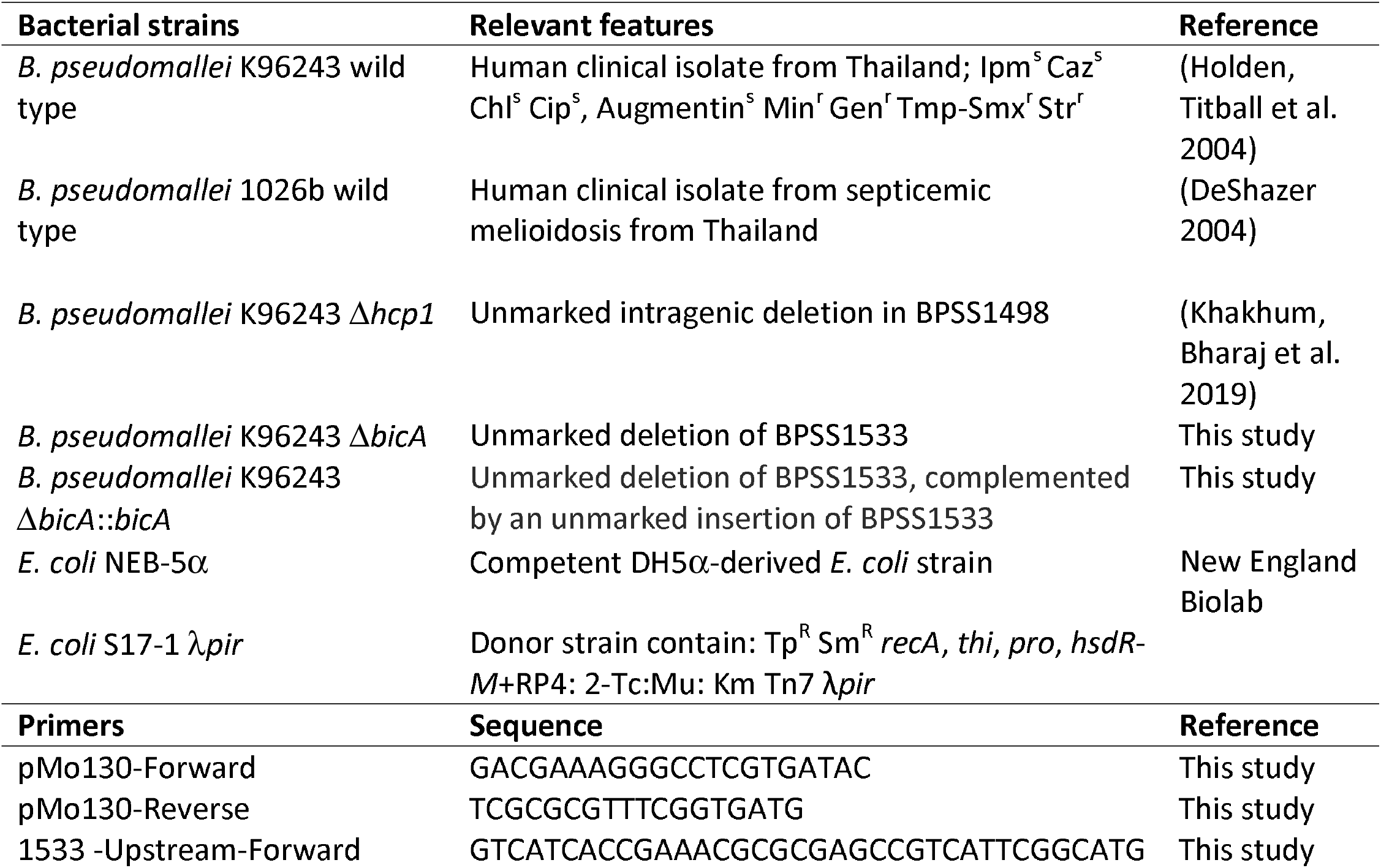
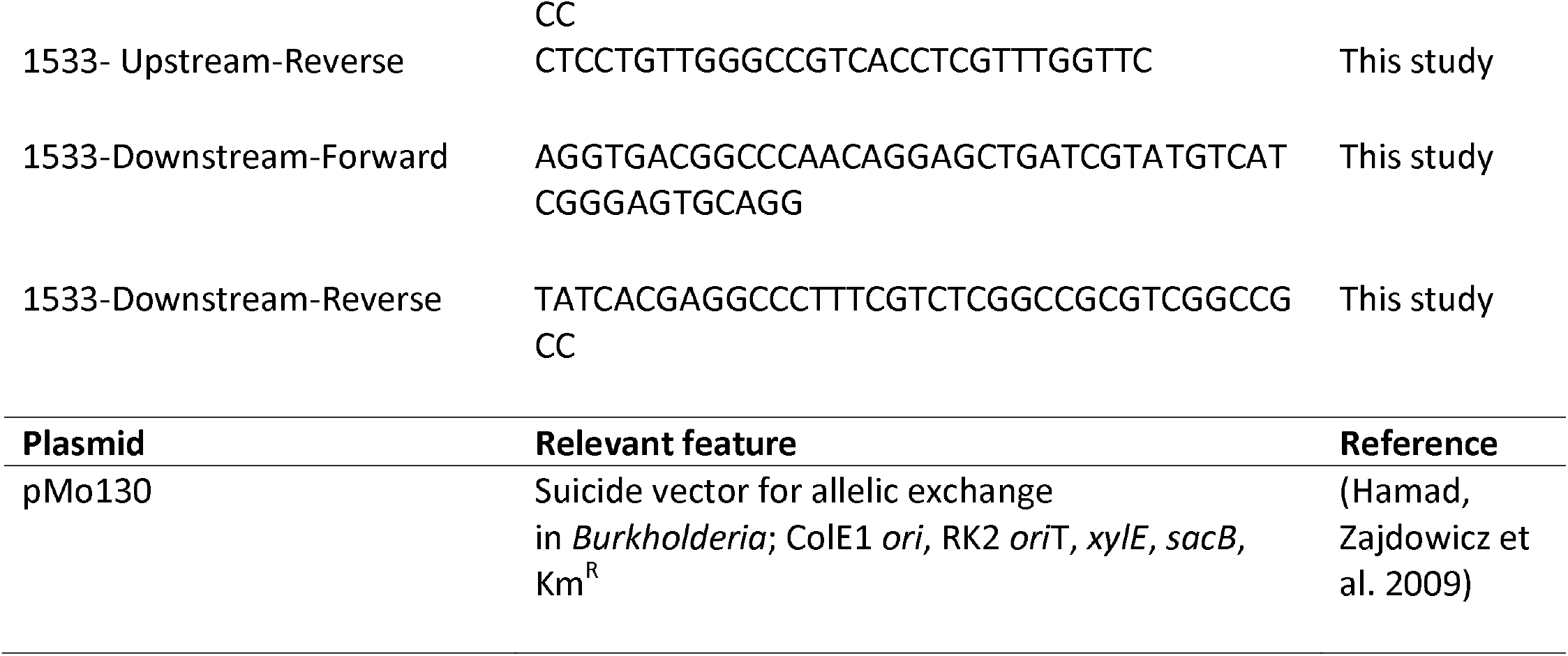
Bacterial strains, primers and plasmid used in this study.

### Transposon library

We used transposon mutants constructed in the prototype *Bpm* strain 1026b. The *Bpm* transposon library was recently acquired from BEI Resources but this library was originally constructed by Dr. Herbert Schweizer, in collaboration with Dr. Brad Borlee and Dr. Colin Manoil (Gallagher, Ramage et al. 2013). This collection of mutants was generated using a comprehensive two-allele sequence-defined transposon mutagenesis of *Bpm* strain 1026b and such library has been used in multiple studies (Borlee, Plumley et al. 2017, Mangalea, Plumley et al. 2017).

### Dual RNA-Seq testing and analysis of the data

#### Bacterial Infection, RNA extraction, and DNase treatment

For RNA-seq analysis, primary IECs were infected as described above with an MOI of 10 for 6 and 12 h with either *Bpm* WT or *Bpm* Δ*hcp1*. After infection, cells were detached, lysed, and total RNA was extracted from whole-cell lysates using Direct-zol RNA Kit (Zymogen, USA) following manufacturer’s instructions. To remove contaminating genomic DNA, samples were treated with 0.25 U of DNase I (Fermentas) per 1 μg of RNA for 45 min at 37 °C. If applicable, RNA quality was checked on the Agilent 2100 Bioanalyzer (Agilent Technologies).

#### rRNA depletion

Bacterial or eukaryotic host rRNA were removed using the RiboMinus™ Human/Mouse Transcriptome Isolation Kit or Bacterial RiboMinus™ Transcriptome Isolation Kit, respectively. Following the manufacturer’s instructions, approx. 500 ng of total, DNase-I-treated RNA from infection samples was used as an input to the ribosomal transcript removal procedure. rRNA-depleted RNA was precipitated in ethanol for 3 h at −20 °C.

#### cDNA library generation and dual RNA Seq analysis

Bacterial and eukaryotic host rRNA was depleted using the RiboMinus™ Human/Mouse and the Bacterial RiboMinus™ Transcriptome Isolation Kits (ThermoFisher, Inc). Following the manufacturer’s instructions, approx. 500 ng of total, DNase-I-treated RNA from infection samples was used as an input to the ribosomal transcript removal procedure. Sequencing libraries were prepared from the depleted RNAs with the TruSeq Stranded RNA Prep Kit (Illumina, Inc) following the manufacturer’s protocol, pooled, and sequenced on a NextSeq 550 High-Output Flow cell using the single-end 75 base protocol.

Host cell reads were mapped and quantified using STAR version 2.7.5c (Dobin, Davis et al. 2013) using the parameters recommended by the ENCODE consortium. The STAR genome index was built with the ENSEMBL Rnor_6.0 reference genome and the version 101 annotation file. Read counts were obtained with the quantMode GeneCounts STAR option. To obtain pathogen read counts the reads were mapped to the *Burkholderia pseudomallei* K96243 reference (GCF_000011545.1) using Bowtie2 version 2.3.4.1 with the –local option (Langmead and Salzberg 2012). Reads per gene were quantified with FeatureCounts, v2.0.1, from the subread software suite (Liao, Smyth et al. 2014), using the version 132 annotation file obtained from the *Burkholderia* Genome Database (Winsor, Khaira et al. 2008). Differential gene expression for the host and pathogen was estimated using the DESeq2 software package, version 1.28.1 following the authors’ vignette (Love, Huber et al. 2014).

### Construction of *bicA* mutant and strain complementation

All cloning methods were performed using the Gibson assembly system and primers used in this study are listed in Table 1. For the mutant construction, upstream (530 bp) and downstream (530 bp) of *Bpm* K96243 of *bicA* gene (BPSS1533) were amplified from genomic DNA and purified before assembly into a linearized PCR product of the suicide vector, pMo130 following the manufacturer’s recommendations. Assembled products were transformed into *E. coli* NEB-5α (NEB, Massachusetts) for clonal screening and the plasmids were confirmed by Sanger sequencing at GENEWIZ. The plasmid containing the fusion of upstream and downstream regions of BPSS153 were ultimately transformed into *E. coli* S17-1 λ*pir* donor strain. Mobilizable vector was introduced into *Bpm* K96243 by biparental mating. Overnight culture (500 µl) of donor *E. coli* S17-1 λ*pir* containing upstream-downstream/pMo130 plasmid and 12 h culture of a recipient *Bpm* K96243 (500 µl) were centrifuged separately or combined with equal volume then resuspended the pellet with 100 µl of PBS. The conjugation mixture was spotted on LB agar and incubated at 37°C for 8 h. Following conjugation, the spots were scraped and resuspended with 1 ml of PBS before plating on LB agar supplemented with 500 µg/ml kanamycin and 30 µg/ml of polymyxin B for *Bpm* transformant selection. Plates were incubated for 48 h at 37°C and the isolated colonies were exposed to 0.45 M pyrocatechol for merodiploid screening. Selected yellow colonies were sub-cultured in LB containing 100 µg/ml of kanamycin then incubated at 37°C with shaking for 12 h. Transformant colonies were grown in YT broth supplemented with 15% sucrose for 4 h at 37°C with shaking followed by serial dilution and plating onto YT agar + 15% sucrose for resolved *Bpm* co-integrant selection. Following 48 h incubation at 37°C, resulting white colonies after exposure to pyrocatechol as mentioned above were analyzed by PCR, and sequenced to confirm gene deletion.

The *in cis* complementation of the *Bpm bicA* mutant was performed by inserting the *bicA* gene back into *Bpm* Δ*bicA* strain using similar manner as the *bicA* mutant construction. Purified PCR amplicon of upstream-BPSS153-downstream was assembled to linearized pMo130 vector PCR product using Gibson assembly then transformed to *E. coli* NEB-5α followed by sub-cloning to *E. coli* S17-1 λ*pir* donor strain. The upstream-BPSS153-downstream/pMo130 plasmid was introduced into *Bpm* Δ*bicA* strain by biparental mating as described above. The clonal selection of complemented *Bpm bicA* mutant was confirmed by PCR and sequencing. Furthermore, the phenotype of both *bicA* mutant and complemented *bicA* was confirmed by plaque formation assay.

### *In vitro* primary epithelial cells infection, plaque formation assay and immunofluorescence microscopic analysis

C57BL/6 mouse primary intestinal epithelial cells (Cell Biologics, Chicago, IL product No. C57-6051), identified by RNAseq results as rat-derived cells were grown in complete primary cell culture medium (Cell Biologics, product No. M6621) following manufacturer’s protocol. Cells were seeded 5 × 10^5^ cells/well in gelatin (Cell Biologics, product No. 6950) coated 24-well culture-plate for survival assay, 1 × 10^6^ cells/well 12-well culture-plate for plaque formation and 2 × 10^5^ cells/well into pre-gelatin coated cover slips in 12-well culture-plate overnight at 37°C, 5% CO_2_ incubator. Cells were infected with *Bpm* K96243 wild type, Δ*bicA* and the complemented Δ*bicA* strain at a multiplicity of infection (MOI) 10 for 1 h, then infected cells were washed twice before incubation with media supplemented with 1 mg/ml kanamycin for an additional 1 h for extracellular bacterial killing. After 1 h, cells were washed and media without antibiotic was replaced then incubated for 3, 6, 12 and 24 h. For survival determination of intracellular bacterial in cells, cells were washed twice and lysed with 0.1% Triton X-100 in PBS prior serial diluted and plated onto LB agar. Plates were incubated at 37°C for 48 h then CFUs were quantified and % intracellular replication was calculated by comparing to input of each bacterial strain. For plaque formation assay, cells at 24 h infection were fixed with 4% paraformaldehyde (PFA) for 30 min and stained with 500 µl Giemsa stain (Gibco) for 30 min. After staining, cells were washed with PBS followed by water before imaging. For immunofluorescence microscopic analysis, after infection, media was removed and replaced with 4% PFA for 30 min. Cells were washed twice with PBS and coverslips were transferred to a new 12-well plate. Fixed cells were lysed with 0.25% Triton X-100 in PBS for 7 min and washed twice with PBS. *Bpm* bacteria were stained with sera collected from mice that immunized with *Bpm* PBK001 live attenuated vaccine at 1:1,000 at RT for 1 h followed by 1:5,000 dilution of goat anti-mouse IgG, IgM, IgA (H+L) secondary antibody conjugated to Alex Flour 488 (Invitrogen) for an additional 1 h. After washing, actin and DNA of cells were stained with 1:10,000 rhodamine phalloidin and DAPI, respectively for 1 h. Coverslips were mounted onto microscopic slides using Prolong diamond antifade mountant (Invitrogen). Slides were visualized with Olympus BX51 upright fluorescence microscope and further analyzed using an ImageJ software, National Institutes of Health (Schindelin, Arganda-Carreras et al. 2012).

### Ethics statement

All manipulations of *Bpm* were conducted in CDC/USDA-approved and registered biosafety level 3 (BSL3) facility at the University of Texas Medical Branch (UTMB) in accordance with approved BSL3 standard operating practices. The animal studies were carried out humanely in strict accordance with the recommendations in the Guide for the Care and Use of Laboratory Animals by the National Institutes of Health. The protocol (IACUC #0503014D) was approved by the Animal Care and Use Committee of UTMB.

### Animal studies

Female 6-to-8-week-old C57BL/6 mice (n = 48) were purchased from Jackson Laboratory (Bar Harbor, ME, USA) and maintained in an Animal Biosafety Level 3 (ABSL3) facility. Animals were housed in microisolator cages under pathogen-free conditions with food and water available *ad libitum* and maintained on a 12 h light cycle.

### Acute (high-dose) and chronic (low-dose) infection models

Food was restricted 12 h before infection but was administered throughout the remainder of the study. For the acute model of infection (n = 8/group), animals were infected with 2.5 × 10^6^ CFU (1 LD_50_) of *Bpm* WT (K96243), *Bpm* Δ*bicA* or *Bpm* Δ*bicA*::*bicA* strain using a plastic oral gavage needle. For the chronic infection model, mice (n = 8/group) were infected with 7.5 × 10^6^ CFU (3 LD_50_). Infected mice were monitored survival and weight for 21- and 35-days post infection. At the study end point, mice were humanely euthanized, and the gastrointestinal tract (stomach, small intestine, colon, and cecum), liver and spleen of surviving animals were collected for bacterial enumeration. All organs were homogenized in 1 mL of PBS, serially diluted, and plated in either LB agar (Liver and spleen) or Ashdown selective medium (GI tract) to quantify bacterial loads.

### Statistical Analysis

All statistical analysis was done using GraphPad Prism software (v8.0). P-values of < 0.05 will be considered statistically significant. Quantitative data is expressed as the mean ± standard error. All data was analyzed for normality before running the corresponding test. Intracellular replication and percent weight change were analyzed by one-way ANOVA followed by Sidak multiple comparison and Kruskal-Wallis test.

## Acknowledgements

This manuscript was partially supported by NIH NIAID grant AI12660101 and UTMB seed funds awarded to AGT. DT was funded by an NIH NIAID Research Supplement for Underrepresented Minorities. JIS-V was supported by the Conacyt ConTex Postdoctoral fellowship. The contents are solely the responsibility of the authors and do not necessarily represent the official views of the NIAID or NIH.

## Author Contributions

JIS-V and DT designed the experiments, JIS-V, DT and NK performed the experiments, SGW analyzed the data. AGT wrote the manuscript. JIS-V, DT, NK, SGW and AGT edited and approved the manuscript.

## Notes

https://www.ncbi.nlm.nih.gov/geo/query/acc.cgi?acc=GSE201263

### Competing Interest Statement

The authors have declared no competing interest.

